# Unraveling the *Myotis* Morass: Ultraconserved-Element Analysis Reveals Introgression, Cryptic Diversity, and Taxonomic Trouble in the Most Species-rich Bat Genus

**DOI:** 10.1101/2022.08.11.503681

**Authors:** Jennifer M. Korstian, Richard D. Stevens, Thomas E. Lee, Robert J. Baker, David A. Ray

## Abstract

Using sequences from 2615 UCE loci and multiple methodologies we inferred phylogenies for the largest genetic dataset of New World *Myotis* to date. The resulting phylogenetic trees were populated with short branch lengths and widespread conflict, hallmarks consistent with rapid adaptive radiations. The degree of conflict observed in *Myotis* has likely contributed to difficulties disentangling deeper evolutionary relationships. Unlike earlier phylogenies based on 1-2 gene sequences, this UCE dataset places *M. brandtii* outside the New World clades. Introgression testing of a small subset of our samples revealed evidence of historical but not contemporary gene flow, suggesting that hybridization occurs less frequently in the Neotropics than the Nearctic. We identified several instances of cryptic lineages within described species as well as several instances of potential taxonomic over-splitting. Evidence from Central and South American localities suggests that diversity in those regions is not fully characterized. In light of the accumulated evidence of the evolutionary complexity in *Myotis* and our survey of the taxonomic implications from our phylogenies it is apparent that the definition of species and regime of species delimitation need to be re-evaluated for *Myotis*. This will require substantial collaboration and sample sharing between geneticists and taxonomists to build a system that is both robust and applicable in a genus as diverse as *Myotis*.

---

Genome sequencing has become a powerful tool for inferring evolutionary relationships among species and reconstructing the tree of life (Hug et al. 2016). However, analyses of these huge datasets have revealed that the traditional tree-like concept may not be best suited to reflect the evolutionary processes involved in creating and maintaining species diversity (Coyne and Orr 2004; Bapteste et al. 2013). Most phylogenetic hypotheses are modeled and viewed as series of simple, bifurcating branches wherein a single parent species gives rise to two independent species that rapidly become distinct and do not interbreed. While situations fitting precisely this definition may occur in nature, such as some island colonization or chromosomal speciation, most speciation events do not occur instantaneously and gene flow between lineages can persist long after the initial divergence (Coyne and Orr 2004; Morales et al. 2017; Morales and Carstens 2018).

Gene flow among lineages and differential selection across the genome make it possible to recover different relationships between lineages depending on the marker(s) used (Prum et al. 2015; Suh et al. 2015; Platt et al. 2018). One way to overcome conflicts among discordant phylogenetic trees is through increasing the number of loci. Such increases allow the resulting species tree to more accurately reflect the genome as a whole (Bejerano et al. 2004; Crawford et al. 2012; Faircloth et al. 2012; McCormack et al. 2012; Prum et al. 2015; Suh et al. 2015; Platt et al. 2018). UCEs (Ultra-Conserved Elements) offer a data rich, sequence-based pool of markers to examine phylogenetic relationships across a wide range of taxa. UCEs are regions of the genome (>200 bp) that are highly conserved across distantly related taxa (Faircloth et al. 2012; McCormack et al. 2012). The sequences from regions surrounding UCEs can be used to infer phylogenies across diverse taxa that may prove challenging or impossible to resolve using other methods.

In bats of the genus *Myotis*, for example, morphological characteristics poorly reflect genetic diversity and species delineations. *Myotis* includes more than 131 species that arose within the last 10-15 million years (Stadelmann et al. 2007; Simmons and Cirranello 2020). Furthermore, several recent genetic studies have found evidence of cryptic diversity within established species boundaries, which may be contributing to the continued phylogenetic uncertainty within the genus (Ruedi et al. 2013; Novaes et al. 2021). A recent study of *M. lucifugus*, for example, found evidence that the current “subspecies” are actually five non-sister species that experience some gene flow between them in regions of secondary contact (Morales and Carstens 2018). Another study examined *Myotis* in the Lesser Antilles and found genetic evidence of a fourth Caribbean endemic species (*M. nyctor*) and evidence of genetically differentiated lineages in the Caribbean and northern South America that may also represent unrecognized species (Larsen et al. 2012a). That study also found that *M. nigricans*, a species widespread throughout South America, could be as many as twelve distinct cryptic lineages. Since then, from within *M. nigricans*, nine new species have been recognized (*M. dinelli, M. clydejonesi, M. bakeri, M. handleyi, M. attenboroughi, M. izecksohni, M. lavali, M. diminutus, M. larensis, M. midastactus*, and *M. nyctor*) (Larsen et al. 2012b; Miranda et al. 2013; Moratelli et al. 2013; Moratelli and Wilson 2014; Moratelli et al. 2016 Dec 15; Barquez et al. 2017; Moratelli et al. 2017; Carrión-Bonilla and Cook 2020; Novaes et al. 2021).

Previous efforts to resolve relationships within *Myotis* using single gene trees and multiple gene trees have resulted in conflicting results (Stadelmann et al. 2007). The strongest phylogenetic hypotheses have therefore been inferred primarily using individual mitochondrial markers. Recently however, Platt et al. (Platt et al. 2018) examined whole mitochondrial genomes and UCEs from 37 individuals (comprising 29 *Myotis* species, primarily from the New World) to determine whether trees constructed with nuclear and mitochondrial markers reflect the same evolutionary history. Using sequences obtained from UCEs, they recovered 294 distinct nuclear UCE trees, all of which differed from those inferred from whole mitochondrial sequences. This observation strongly suggests that the mitochondrial and nuclear genomes experienced different evolutionary histories. Furthermore, half of the individual gene trees generated from individual UCE loci were inconsistent with the consensus species tree, likely due to incomplete lineage sorting (ILS), hybridization, and/or phylogenetic error. Despite this, one single nuclear phylogeny was recovered from the UCE data in almost 25% of the trees. By contrast, the mitogenome trees recovered a different phylogeny and did not have a single tree that was recovered with such high frequency. This is all rather problematic because inferences based on both mitochondrial and nuclear trees have been used to inform conservation efforts and to support evolutionary and population genetic hypotheses (Forest et al. 2015; Rubinoff et al. 2020; Grimshaw et al. 2022).

A weakness of Platt et al.’s work was the relatively limited sampling. Most species were represented by only one individual and relatively few New World *Myotis* were represented out of the over 100 *Myotis* species that likely exist. Increased sampling can help resolve ambiguous relationships and identify the causes behind such ambiguity. Therefore, we have expanded sampling by >10X to investigate the relationships among New World *Myotis* and to reveal patterns of ILS, hybridization, and possible cryptic speciation in the clade. We also incorporate the new species mentioned above into a single large-scale phylogenic inference, reconcile existing species designations with our UCE phylogeny, and assess the number of cryptic lineages requiring further investigation.

## Methods

### Taxon Selection

We targeted representatives of as many named species of *Myotis* in the New World as possible as well as several Old World *Myotis* and non-*Myotis* vespertilionid bats to act as outgroups (Fig. 1a & 1b). A detailed table of all samples can be found in Supplemental Material. When possible, we included a minimum of two individuals for every New World *Myotis* species, particularly when those species or species complexes are associated with large ranges or potential cryptic lineages (e.g. *M. nigricans, M. albescens, M. martiniquensis/nyctor*) (Larsen et al. 2012a; Platt et al. 2018). This project relies heavily on tissue and DNA samples from museum research collections. The majority of our samples were obtained from the Natural Sciences Research Laboratory at the Museum of Texas Tech University (TTUNSRL). Additional samples were loaned from the Museum of Southwestern Biology (MSB), the Abilene Cristian University Natural History Collection (ACUNHC), the Angelo State Natural History Collection (ASNHC), the Sam Noble Oklahoma Museum of Natural History (SNOMNH), the Museum of Vertebrate Zoology (MVZ), the American Museum of Natural History (AMNH), the National Museum of Natural History (NMNH), the Louisiana Museum of Natural History (LSUMZ), as well as private research collections. For samples with GPS coordinates or detailed locality descriptions that allowed us to obtain approximate coordinates using Google Maps, we mapped the collection localities using the ggplot R package (Wickhman 2016).

**Figure 1.**
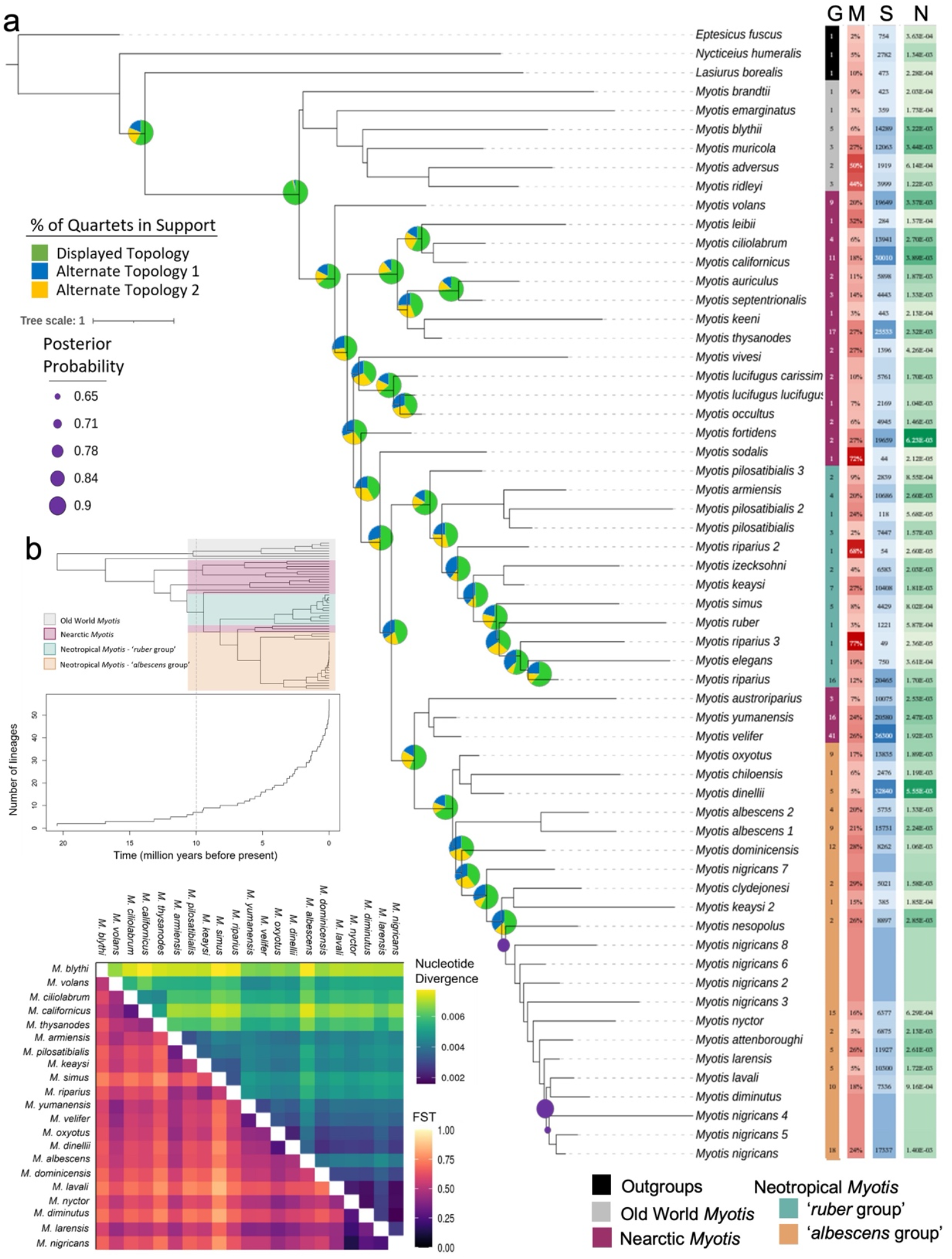
a) *Myotis* ASTRAL_S species tree inferred from 2,615 UCE gene trees. Pie charts show the percentage of trees supporting each of the three possible quartet topologies. All branches received posterior probabilities > 0.90 unless indicated with purple dots. Colored bars on the right of the tree show the geographic origins of samples (labeled G, key bottom right), percent missing data (M), number of segregating sites (S), and nuclotide diversity (N). Note: all *M. nigricans* lineages were pooled for calculating descriptive statistics. b) Top: Time calibrated phylogenetic tree of *Myotis* lineages in our UCE dataset. Highlighting indicates phylogeographic membership of the lineages as in part a. Bottom: Plot of the number of *Myotis* lineages over time. The grey dotted line corresponds to 10 MYA in both figures. c) Heatmap of pairwise FST and nucleotide diversity values for species with at least 4 individuals sampled. Nucleotide divergences are shown above the diagonal and FST is shown below the diagonal.

### DNA Extraction, Library Preparation, and Sequencing

We obtained liver, muscle, or wing punch tissue samples for 341 bats. DNA was isolated using the Qiagen DNEasy extraction kit (Qiagen, Germantown, MD) using the manufacturers protocol. UCE loci were captured as described in Platt et al. (2018). Briefly, this procedure captures UCE loci from fragmented genomic libraries using the Tetrapods 5K version 1 baits from MYcroarray (Ann Arbor, MI, USA). Libraries were sequenced at the EHS DNA Laboratory at the University of Georgia, producing 151 bp paired-end reads. In addition to sequences produced from tissue samples, we included previously published UCE reads by Platt et al. (2018).

### UCE Processing

After demultiplexing, all samples and reads were trimmed to remove adapter sequences and low-quality reads using Trimmomatic (Bolger et al. 2014) and assembled into contigs using Spades (Grabherr et al. 2011). UCE sequences were processed with the PHYLUCE (Faircloth 2016) pipeline where we retained the default values unless otherwise specified. In brief, the contigs of the assembled reads were matched to the PHYLUCE’s UCE-5k-probe set (Faircloth et al. 2012). Regions matching probes plus 500bp of flanking sequence were then extracted from the contigs, alignments were produced for each locus using MAFFT (Katoh and Standley 2013), and edge trimmed.

The sequences in these alignments were then used as a template for each individual in the phasing workflow from the PHYLUCE pipeline. In this workflow, the reads were mapped to the consensus sequence from that individual and two allele fasta files were obtained. Those allele fasta files were treated as the new ‘assembled contigs’ in a further round of processing and were matched to contigs, extracted, aligned by locus, and edge trimmed as before. Individuals with fewer than 1000 UCE loci were dropped from the analysis and only alignments containing sequences for at least 75% of taxa were retained for further analysis. For each individual retained in the analysis, we included both of the phased allele sequences, essentially doubling our sample size.

### UCE Phylogeny

We inferred individual gene-trees for each UCE locus with RAxML-NG (Kozlov et al. 2019) with a single analysis that implements a GTR+G model and 100 bootstrap trees after selecting the tree with the best maximum likelihood (ML) from the 10 parsimony-based starting tree topologies. In addition to a single best ML tree, RAxML-NG also produces a tree where near-zero branches are collapsed. We used the collapsed versions of the best ML trees to infer the species trees using ASTRAL-MP (Sayyari and Mirarab 2018; Zhang et al. 2018).

The ASTRAL analysis was run in two ways: 1) each allele sequence was assumed to be its own ‘species’, 2) all allele sequences were assigned to species based on their updated taxonomic IDs (see Species Identification) and a species tree was inferred from the gene trees. We ran a second analysis to score the species trees and produce detailed annotations for each branch. The resulting trees were visualized using the Interactive Tree Of Life (iTOL)(Letunic and Bork 2021) and combined with supplementary graphics to depict phylogenetic uncertainty.

### Species Identification

Identification of our samples was an enormous challenge because of the high degree of morphological similarity within the genus, recent revisions to taxonomy, inaccurate identification of museum vouchers, inconsistencies in taxonomic classifications, and a lack of clear consensus as to what constitutes a species. Defining/redefining species delimitation within *Myotis* was beyond the scope of this project because we were able to physically examine only a small proportion of the samples in this study. This meant that many of our identifications are inferred by genetic similarity alone and they should ideally be confirmed with morphology. Therefore, we are treating our species identifications as hypotheses that will need to be compared to the existing species to identify problematic taxa that merit further evaluation. We defined a species as a monophyletic population that is genetically distinct and with unique diagnostic morphological characteristics. While most of the currently named species fit well within this definition, we found several instances where putative species do not meet these criteria. To avoid confusion and prevent this manuscript from becoming a large-scale taxonomic revision, we have preserved species that are currently recognized by the taxonomic literature even if they do not meet those criteria (i.e. we did not collapse/subsume a named species into another).

We used an iterative process to obtain final species IDs. First, we treated each sample as its own ‘species’ for ASTRAL analysis, a process referred to as ‘individual mode’, to produce a draft UCE tree and group samples based on topology and the initial species designations (when available). Samples forming monophyletic groups with posterior probabilities greater than 0.8 were grouped together as hypothesized species. We then examined the 37 museum vouchers present at the TTUNSRL to confirm the genetic species identifications for those samples identified only to the genus or misidentified. Several samples had been examined in prior studies; for those cases we used the most recently published identification. Vouchers were identified using (Moratelli et al. 2013; York et al. 2019) as guides, in conjunction with descriptions of new species that have been named more recently (Larsen et al. 2012b; Miranda et al. 2013; Moratelli et al. 2013; Moratelli and Wilson 2014; Moratelli et al. 2016 Dec 15; Barquez et al. 2017; Moratelli et al. 2017; Moratelli et al. 2019; Carrión-Bonilla and Cook 2020; Novaes et al. 2021).

After initial identifications, we used ASTRAL to make a new tree in ‘species mode’ wherein all samples belonging to each of our designated lineages are treated as a single unit that cannot be split. To minimize the likelihood of including a sample in the wrong group, we used a conservative approach by maintaining several smaller lineages over one combined, well-diverged lineage when possible because the ASTRAL analysis does not allow a single ‘species’ to be split but will allow several ‘species’ to be combined later. We tested the resulting ‘species’ tree to determine if a branch should be replaced with a polytomy because of insufficient support. This test allowed us to identify instances where several lineages should be combined. Except in instances when doing so would combine two currently accepted species, we collapsed branches and repeated as needed to eliminate poorly supported divisions between lineages as indicated by the polytomy testing or ASTRAL support values. Finally, we used FST and nucleotide diversities to identify instances where we had been maintaining multiple lineages with exceptionally low pairwise nucleotide divergences and FST values, a pattern that indicates that those lineages should be combined and reanalyzed. After the completion of this process, we considered each surviving group to be a species. For simplicity and clarity, during our discussion of our ASTRAL UCE trees, we will refer to those created in ‘species mode’ as ASTRAL_S and those created in ‘individual mode’ as ASTRAL_I and have only shown one allele per individual in the ASTRAL_I trees. The full ASTRAL_I tree showing both alleles for all individuals can be found in Supplemental Material.

We calculated descriptive statistics on each species to characterize the diversity within each group. We used PHYLUCE to concatenate all of the UCE alignments by sample and then calculated nucleotide diversity (pi), haplotype diversity, number of segregating sites, and pairwise FST values for each species using the PopGenome R package (Pfeifer et al. 2014). We created heatmaps of pairwise FST and nucleotide divergence values for all species with at least four individuals using the ggplot R package (Wickhman 2016). A neighbor-joining tree was inferred from the concatenated alignments and a principal component analysis was conducted using the adegenet R package (Jombart and Ahmed 2011).

### Cytochrome b Trees

Because many of the samples utilized in this study had previously been sequenced for cytochrome b (cytb)-based phylogenies, we downloaded existing cytb sequences whenever possible (n=95). Accession numbers of all cytb sequences used in this study can be found in Supplemental Material. We were able to obtain sequences for an additional 50 individuals by taking advantage of non-targeted mitochondrial sequences in the reads produced for our UCE sequencing. We used Trinity (Grabherr et al. 2011) to assemble the reads for each sample using a custom library of nine cytb reference sequences obtained from GenBank for our species of interest (fasta library available on Dryad). We used Muscle (Edgar 2004) to align the cytb sequences and trimmed all sequences to 1140 bp. We inferred a tree using RAxML-NG (Kozlov et al. 2019) with a single analysis that implements a GTR+G mutation model and 100 bootstrap trees after selecting the tree with the best maximum likelihood (ML) from the 10 parsimony-based starting tree topologies. The resulting trees were visualized using the Interactive Tree Of Life (iTOL)(Letunic and Bork 2021).

### Testing for Gene Flow

With a large phylogeny, it is difficult to explore all possible pathways for signs of introgression because the number of pathways grows exponentially such that to examine our entire phased dataset would require examining nearly 4 billion quartets. To reduce the search space for introgression testing, we constructed a reduced dataset consisting of concatenated UCE sequences for 12 species, each represented by a single allele from a single individual per species. By limiting our introgression testing to 12 species, we were able to reduce the search space to just 495 quartets. Reducing our dataset and targeting specific species allowed us to test several hypotheses about gene flow within *Myotis*.

For the 12 species, we selected *M. emarginatus*, the most closely related Old World species in our dataset (according to our UCE tree) with a current distribution that reduces the possibility of gene flow with New World species, to serve as the outgroup for these tests. *M. ciliolabrum* and *M. californicus* were included because hybridization is one possible contributor to their high genetic similarity (Rodriguez and Ammerman 2004). *M. nigricans, M. albescens* and *M. oxyotus* were included as representatives of the neotropical ‘*albescens* group’ and *M. velifer* was selected as the Nearctic representative from that clade (Moratelli et al. 2013). *M. riparius* and *M. pilosatibialis* were selected for the neotropical ‘*ruber* group’ (Moratelli et al. 2013). *M. lucifugus* was included because it is known to hybridize (Morales and Carstens 2018). *M. brandtii* was included because evidence based on SINE insertions (Korstian et al. 2022) indicated historical gene flow between it and several North American *Myotis* species. Finally, *M. volans* was selected because of its position sister to all other New World species in our ASTRAL_S UCE-species tree (Fig. 1a), as we hypothesized that historical gene flow could drive taxonomic uncertainty within the tree that could result in such a placement.

The quartet asymmetry test we employed makes use of an ASTRAL tree to determine the “accepted” topology. Though designed for use with retrophylogenomic data (Springer et al. 2020), we have utilized concatenated UCE sequences instead and applied the method to SNPs. Since retrotransposons are less prone to homoplasy than SNPs and SNPs can have up to four possible states instead of the binary presence/absence states of retrotransposons, the relative efficacy of this strategy with sequence-based data has not been thoroughly tested. Despite the potential shortcomings of this analysis strategy, we believe this test can still provide general insights into the overall prevalence of gene flow within the subset of samples we tested. For each of the 495 possible quartets formed from our 12 species, we used PAUP* (Swofford 2002) and custom scripts to score the three possible tree topologies.

Additional details on the custom scripts and the scoring process can be found in (Korstian et al. 2022). Using those tree scores, we calculated the number of characters in support of each topology and compared the observed frequencies to the expectations under the influence of ILS alone (i.e. that the two alternate topologies will receive roughly equal support) using a χ^2^ test. We then applied a Holm-Bonferroni correction for multiple tests (Holm 1979) using an Excel calculator designed for the purpose (Gaetano 2018). We then grouped the quartets indicating introgression such that the fewest possible introgression events were required to obtain that pattern. In other words, when many quartets supported introgression between species A and species B and between species A and C, we concluded that there was a single introgression pathway between A and the shared common ancestor of B and C, rather than two separate pathways.

### Lineages Through Time

To investigate the rate of speciation in *Myotis*, we created a time-calibrated version of our UCE species tree using the penalized likelihood method implemented in the *chronos* module of the ape R package (Paradis E 2019). Calibration date minimum and maximum values for several key nodes were obtained from (Ruedi et al. 2013). We employed a relaxed likelihood model and a lambda of 1. Lineage through time plots were created using the *lttplot* module of the ape R package.

## Results

### Summary Statistics

We recovered phased UCE sequence data for 341 individuals of which 278 passed all quality control measures and were utilized for this analysis. Ten samples were identified as non-target species and were excluded from the study, four samples were excluded because of potential sample mix-ups, and 49 were excluded for not having sequences for at least 1000 UCE loci. Our dataset incorporates representatives of at least 37 of the 52 New World *Myotis* species currently recognized (Simmons and Cirranello 2020). With these data we were able to determine species level identifications for approximately 70 samples that had been previously misidentified or only identified to genus. The number of individuals per species varied widely (range: 1-41) with an average of 5.8 ± 7. 3 (Fig. 1a).

The final ASTRAL_S phylogeny (Fig. 1a) was constructed from 2615 UCE alignments with sequences present for at least 75% of the samples. Because both alleles were assembled for each individual, the effective number of samples was 556. UCE alignments contained an average of 510 individuals per locus (range: 418-556). When concatenated by individual, the total length of all loci was 2,078,858 bp yielding 476,245 informative sites (average=182 per locus). The average sequence length per locus was 795 bp (range: 315-2066 bp).

The average nucleotide divergence between all species pairs in our study was 0.0056 (± 0.0034) substitutions per site between species. Nucleotide divergences between all *Myotis* species were lower, with an average of 0.0048 (± 0.0021) substitutions per site. Within the New World, the average nucleotide divergence between any two *Myotis* was 0.0046 (± 0.0021) substitutions per site. Average nucleotide divergences within each species (Fig. 1c) varied widely with an average value of 1.56×10^−3^ substitutions per site (range: 2.12×10^−5^ – 6.23×10^−3^). The average FST between all species pairs in our dataset was 0.66 (± 0.23, Fig. 3.2). Within all *Myotis* the average FST value was 0.63 (± 0.23) and within just the New World *Myotis* FST was 0.59 (± 0.23).

**Figure 2.**
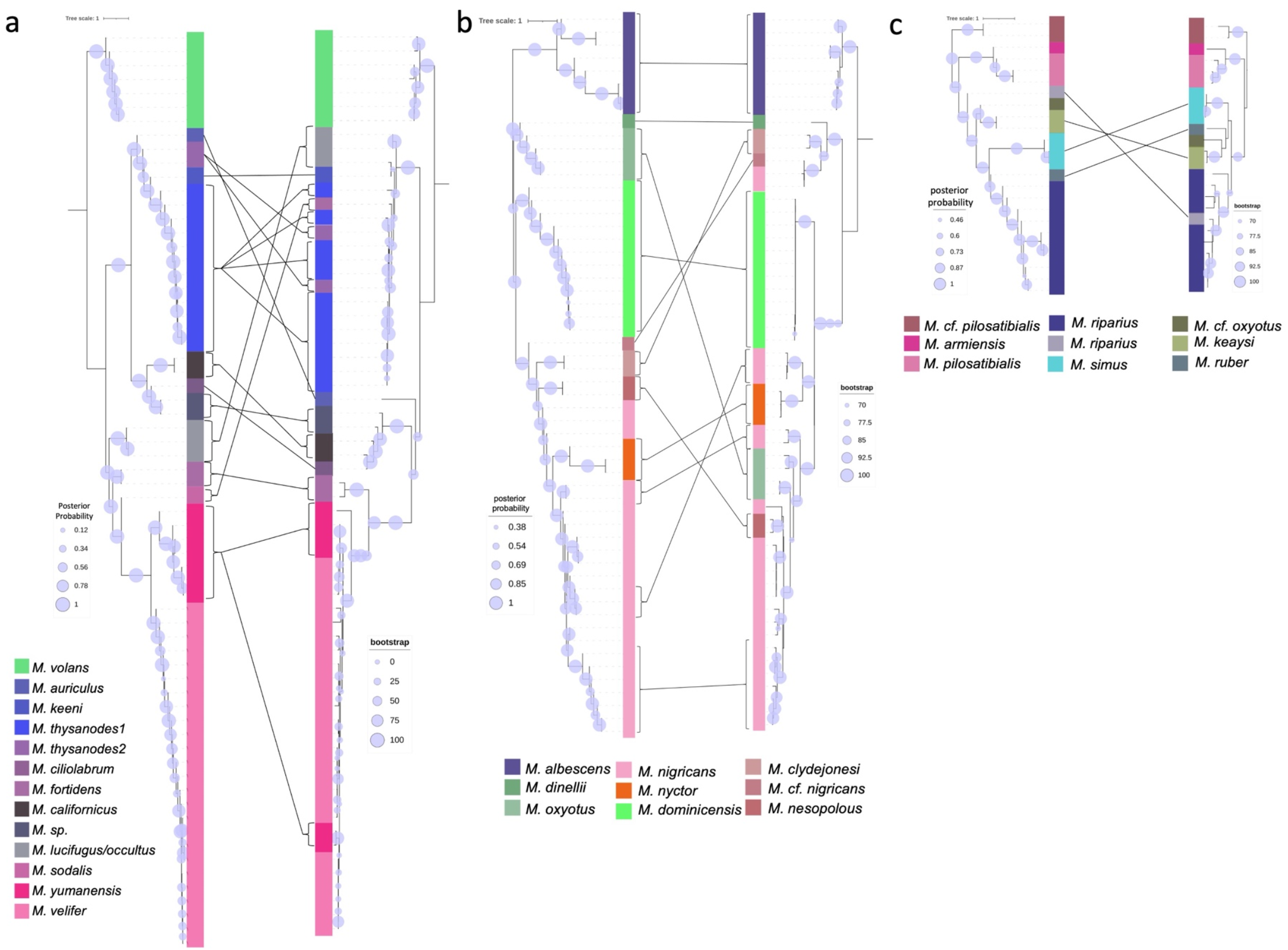
Comparison of tree conflict by marker for the three broad phylogeographic *Myotis* groups. a) Nearctic group, b) neotropical ‘*albescens* group’, c) neotropical ‘*ruber* group’ This is a subset of the individuals shown in Fig. 1a. For each group, the left tree displays the ASTRAL_I UCE phylogeny while the right displays the Maximum Likelihood Cytochrome-b phylogeny. The same individuals are shown in each tree pair. Lines between trees denote notable topological differences between the markers. Colors for highlighting by species are an approximated average color for that species as determined by the first three components of the PCAs shown in Supplemental Material.

**Figure 3.**
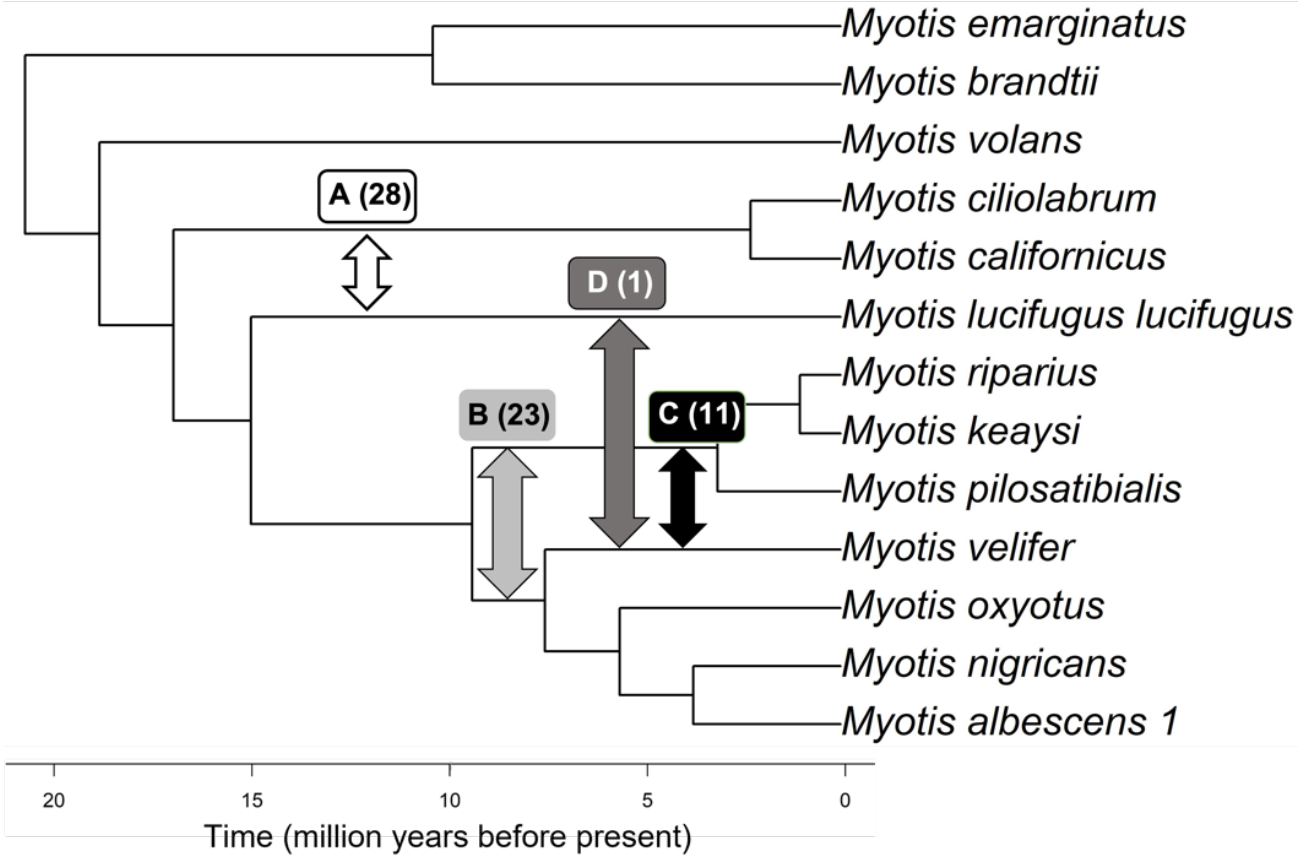
Proposed Introgression Pathways identified using quartet asymmetry tests. 63 quartets indicated significant deviation from expectations under ILS. Pathways are identified by letters with number of quartets supporting in parentheses. Branch lengths are time calibrated as in Fig. 1a.

As seen in other studies, there are a number of species that do not follow established biogeographic trends. In particular, three Nearctic species (*M. yumanensis, M. velifer*, and *M. austroriparius*) were found basal to the ‘*albescens* group’ of neotropical bats (Stadelmann et al. 2007; Agnarsson et al. 2011; Ruedi et al. 2013; Platt et al. 2018) (Fig. 1a). This tree topology is supported by more than half of the quartets in our ASTRAL_S analysis which suggests that the ‘albescens group’ of bats likely evolved from an ancestor of Nearctic species that colonized the Neotropics approximately 5 MYA (Fig. 1b).

Notably, support for the branch separating the Old World and New World *Myotis* is nearly universal (Illustrated by the pie charts in Fig. 1a) which makes this the first phylogenetic study to place *M. brandtii* within the Old World clade as opposed to within the New World (Stadelmann et al. 2007; Agnarsson et al. 2011; Ruedi et al. 2013; Platt et al. 2018). This is consistent with its current distribution in Europe and Asia (Seim et al. 2013).

The ASTRAL_S tree is composed primarily of many, very short branches (Fig. 1a). The longest non-terminal branch within *Myotis* was 1.5 coalescent units (CU) but the average was much smaller at 0.32 CU. These short branch lengths are consistent with expectations for species that experienced extremely rapid radiations. The lineages through time plot (Fig. 1b) shows a higher rate of new lineages created within the last 10 MYA (50 lineages) than in the 10 million years preceding it (7 lineages). Furthermore, the short branch lengths that result from such rapid diversification indicate that the probability of ILS in *Myotis* is high. The quartet support values for key branches of the inferred ASTRAL_S phylogeny (Fig. 1a) highlight the extent of phylogenetic conflict within this genus. For many branches, the three possible quartet topologies were supported with roughly equal frequency (Fig. 1a).

For individuals with both UCE and cytb sequences available, the inferred phylogenies for both markers have been split into each of the three major phylogeographic groups for ease of visualization. The UCE trees and the cytb trees do not recover identical topologies for any of those clades (Fig. 2). While many of the differences between trees are relatively minor, some suggest vastly different evolutionary relationships and population structure. A detailed examination of our results for each of the three phylogeographic groups is included in the Supplemental Material.

### Introgression Testing

Of the 495 possible quartets in our reduced 12 sample dataset, 63 showed significant deviations from expectations under ILS alone, suggesting gene flow is occurring in the quartet (Fig. 3). We have grouped these quartets into the four most parsimonious potential introgression pathways (Fig. 3). There were several notable instances where gene flow was not detected. We found no evidence of gene flow occurring between *M. ciliolabrum* or *M. californicus*. Nor was there evidence of gene flow among species belonging to the same neotropical clade. For example, no quartets involving introgression between *M. nigricans* and *M. albescens* were detected. The same was also true for *M. pilosatibialis* and *M. riparius*. In contrast, 11 quartets support gene flow between the common ancestor of the *M. riparius* clade and the common ancestor of the clade containing *M. nigricans* and *M. oxyotus* and an additional 23 to *M. velifer*. We found 28 quartets that support gene flow between *M. lucifugus* and the common ancestor of *M. californicus* and *M. ciliolabrum*. Three of the four potential gene flow pathways and 40 of the quartets we examined involve at least one Nearctic species. Only 23 quartets support gene flow within Neotropical species only, suggesting the possibility that gene flow may be more common among Nearctic species than those in the Neotropics.

Based on our time-calibrated phylogeny (Fig. 1b), we estimate that the introgression we detected occurred from approximately 4-15 MYBP and is indicative of historical gene flow associated with range expansions and vicariance events as opposed to contemporary gene flow. It is highly probable that increasing the search space to include more species and quartets will identify additional evidence of gene flow between lineages and extant species within this genus.

## Discussion

The *Myotis* phylogenies we produced clearly indicate that phylogenetic conflict is widespread within the genus. Traditionally, evolutionary relationships are represented in terms of a strictly bifurcating tree wherein one parent species gives rise to two offspring species. Under that model, portions of a tree that cannot resolve each evolutionary event into a bifurcation would have been considered as flawed. Genomic data has provided numerous opportunities to see that speciation does not always follow a strictly bifurcating modality. With this knowledge comes the understanding that a phylogenetic tree populated with short branch lengths and high conflict has not necessarily failed in uncovering the evolutionary relationships among its leaves. In other words, these regions are not ‘bugs’ they are ‘features’ that can provide additional information about the evolutionary dynamics of a speciation event that could not otherwise be seen in a strictly bifurcating model. Indeed, the short branch lengths and high conflict are consistent with radiations so rapid that the differentiation in many in *Myotis* may be effectively viewed as instantaneous.

This degree of conflict observed in *Myotis* has likely contributed to difficulty disentangling the deeper evolutionary relationships in the past. This tree and the TE-based tree presented in Korstian et al. (2022) are the first to place *M. brandtii* outside the New World clades. Many of the previous analyses were based on mitochondrial DNA. Topologies resulting from the use of mitochondrial DNA to infer phylogenetic relationships can reflect only the evolutionary trajectory experienced by the mitochondria, which is a single uniparentally inherited, non-recombining locus. By employing multiple recombining loci (such as UCEs) to infer relationships, one can construct phylogenies that reflect the evolutionary history of the genome as opposed to that of a single organelle or gene. Both the number of loci examined and the locations of those loci within the genome may play a critical role in determining if a phylogenic inference method can overcome the confounding effects of ILS and introgression and provide a more accurate tree.

Rodriguez and Ammerman (2004) found that morphology-based species designations are inconsistent with mitochondrial DNA divergence for *M. ciliolabrum*, and *M. californicus*. We hypothesized that gene flow between *M. ciliolabrum* and *M. californicus* might be a contributing factor in the ongoing difficulty with identification and delineation between the two species, but our introgression testing found no evidence in support of that hypothesis. Instead, the absence of introgression would be a compelling argument in favor of maintaining *M. ciliolabrum* and *M. californicus* as two species, rather than a single, morphologically variable species, both possible solutions suggested by Rodriguez and Ammerman (Rodriguez and Ammerman 2004). Though it should be noted that the introgression tests utilized a single representative for each species, particularly with taxa that are frequently misidentified. Sampling with multiple individuals across the entire distribution of both species would be ideal to confirm the absence of introgression between *M. ciliolabrum* and *M. californicus*.

This UCE-based tree is also consistent with the dispersal pathways proposed for the arrival of *Myotis* in the New World by Ruedi et al. (Ruedi et al. 2013) based on one mitochondrial gene (cytochrome b: Cyt-*b*) and one nuclear gene (Recombination Activating Gene 2: RAG2). Their proposed pathways have *Myotis* arriving in the New World from the eastern palearctic around 12 MYA (Ruedi et al. 2013). Subsequent radiations split the New World clade into two primary subclades roughly corresponding to geography (Neotropical and Nearctic subclades) and one more usual clade that includes two species (*M. brandtii* and *M. gracilis*) from the Palearctic/Asia (Ruedi et al. 2013). Given evidence of introgression between *M. velifer* and the *ruber* group and the basal position of *M. velifer* relative to the *albescens* group, we hypothesize that common ancestor of *M. velifer, M. yumanensis*, and *M. austroriparius* was one of the first of this genus to expand into the neotropics and that during that period geneflow between incipient species occurred more frequently than in current populations.

In contrast with the extensive gene flow identified by a prior study using transposable elements (Korstian et al. 2022), we did not find evidence of contemporary hybridization among extant species. Indeed, only one quartet identified gene flow between a single species pair (*M. lucifugus* and *M. velifer*), though had additional species from the Nearctic been included, we may have observed different results. Our tests revealed strong support (23 quartets) for historical hybridization between the two main clades of *Myotis* in the neotropics, the ‘*albescens* group’ and the ‘*ruber* group’. An additional 11 quartets supported gene flow between *M. velifer* and the common ancestor of *M. riparius* and *M. pilosatibialis*. Despite the evidence of historical gene flow, we found no evidence of contemporary gene flow within the ‘*albescens* group’ or the ‘*ruber* group’. This suggests the possibility that *Myotis* populations in the Neotropics hybridize less frequently than those in the Nearctic. Bats in the Nearctic often employ large scale migratory behavior and often occupy large ranges (Davis and Hitchcock 1965; Winhold and Kurta 2006; White et al. 2017) whereas those in the Neotropics move on a smaller scale and species appear to have smaller ranges with patchy distributions (Esbérard et al. 2017). These factors suggest that migratory behavior and the introgression accompanying it may result in homogenization across Nearctic lineages while the patchy ranges and lack of migration may have accelerated speciation in the Neotropics. This may contribute to why the number of *Myotis* species found in the Neotropics is greater than that found in the Nearctic despite having more recently radiated into the Neotropics.

Evidence from the few Central American localities in this study suggests that *Myotis* diversity within Central America is not fully characterized. The geographic sampling for this study was by necessity opportunistic and has several large voids as a result. Notably, Brazil and Chile are unsampled while Mexico and Central America are underrepresented and are likely to have similar patterns of local adaptation and undiagnosed diversity as the other neotropical species examined in this study. In addition, given the sheer number of cryptic species being found in the neotropics, it seems likely that the same will be true of other tropical regions which have an even poorer record of sampling than the Americas have.

In light of the accumulated evidence of the evolutionary complexity in *Myotis*, it is apparent that the definition of species and regime of species delimitation need to be re-evaluated for *Myotis*. Any such system would require extensive re-examination using both morphological and genomic data and would need to be capable of incorporating gene flow among some extant species. To that end, it will be necessary to build a more concrete set of guidelines about what constitutes a species within this genus. This will require more collaboration and sample sharing between geneticists and taxonomists to build a system that is both robust and applicable in a genus as diverse as *Myotis*. DeRaad et al. (2022) described a strategy using species trees, species delimitation, demographic model testing, and tests for gene flow to clarify complex speciation signatures in scrub-jays. Multi-faceted genomic strategies such as these may provide potential answers within *Myotis* as well.

The UCE phylogeny we produced is one of the first to examine the implications of gene flow and incomplete lineage sorting within *Myotis*. Additional investigation will yield further insights into the extent and frequency of hybridization within the genus. Ideally, these investigations will continue with UCEs and make use of a universal probe set (i.e. the UCE-5k probes used in this project) so that the subsequent analyses can build upon this dataset. This strategy would allow some or all of the present dataset to be reanalyzed as more data becomes available and as taxonomic revisions are undertaken, much as cytb sequences were used previously. Throughout this project, we have identified instances where additional sampling from across the entire distribution of a species would aid in answering unresolved questions. In light of the evidence of gene flow presented in this study and in other recent studies, much is yet to be discovered about the extent of contemporary gene flow between Nearctic species.

Understanding the evolutionary relationships and dynamics of species is vital to interpreting data and implementing management decisions. For instance, a conservation plan for a single widespread species will be different from the strategy developed if that single species is in fact two species (a mainland species and an island endemic species). This work underscores the limitations of the current species delineations within *Myotis* and provides a framework for building upon our current understanding of interspecies relationships and boundaries within the Americas. With bat populations constantly under pressure from anthropogenic threats, it is more important than ever to have a firm understanding of what biodiversity exists within a genus as well as what factors have shaped that diversity in order to best determine how to preserve it.

## Supplementary Material

Data available from the Dryad Digital Repository (Repository In progress)

## Acknowledgements

This study would not have been possible without the contributions of Robert J. Baker who conceived and began the project prior to his passing. This research was funded by a Horn Professorship (RJB), the Clark Stevens Endowed Professorship (TEL), and the National Science Foundation, grant numbers 2032006, 1838283, and 1355176 and by the Texas Tech University College of Arts and Sciences. Heath Garner and the Natural Science Research Laboratory of the Museum of Texas Tech University provided valuable tissue samples and logistical support. The TTU High Performance Computing Cluster provided computational resources and technical support. Library preparation and sequencing was completed by the Environmental Health Science DNA Lab at the University of Georgia. Matt Johnson contributed advice and computational support. Thank you to Francisco Castellanos for help with identifications. Thanks to Heath Garner for logistical support. Thank you to Museum of Southwestern Biology (MSB), the Angelo State Natural History Collection (ASNHC), the Abilene Cristian University Natural History Collection (ACUNHC), the Sam Noble Oklahoma Museum of Natural History (SNOMNH), the Museum of Vertebrate Zoology (MVZ), the American Museum of Natural History (AMNH), the National Museum of Natural History (NMNH), and the Louisiana Museum of Natural History (LSUMZ) for providing tissue samples.

## Data Availability

Raw UCE read data for all samples is publicly available via NCBI Sequence Read Archive under the BioProject ID PRJNA834103. Cytb sequences generated by this project are publicly available via NCBI GenBank BioProject ID (PENDING)

Alignments (UCE and cytb) are available for download at (DRYAD info pending).

